# Discrete Inverse Rendering: Biological Image Analysis with Integer Programming

**DOI:** 10.64898/2026.07.23.740284

**Authors:** Frans Zdyb, Julius B. Kirkegaard

**Author notes:** Corresponding authors, &.

## Abstract

Biological image and video analysis is full of discrete decisions: whether an object is present, which multi-hypothesis detections are real, whether two detections are tracking the same object, or whether a cell divides or not. Standard pipelines resolve these locally and in sequence, e.g through non-max suppression, per-frame segmentation, distance-based linking, or dedicated lineage rules. Image evidence and temporal evidence are thus rarely weighed against each other in a single objective. We recast these discrete steps as *discrete inverse rendering* : candidate renderings are generated, and then a integer programming solver selects the subset that best reconstructs the observed video subject to problem-specific constraints. We demonstrate that the same solver and the same reconstruction principle can handle three otherwise separate motifs: suppression of overlapping detections in dense *C. elegans* tracking, extraction of a single connected curve in sperm tracking, and event-structured tracking in single cell tracking.

## 1 Introduction

Inverse rendering provides a natural framework for many tasks in image analysis. Instead of treating an image as a collection of pixels to be classified, inverse rendering asks what configuration of objects could have generated the observed image or video. In biology, these objects may be cells, nuclei, splines, flagella, filaments, membranes, or other shape primitives.

Many object properties are continuous: position, shape, intensity, curvature, or deformation. But many of the most important decisions are discrete. A candidate object is either present or absent. Two overlapping hypotheses either represent the same object or two different objects. A curve fragment either belongs to the flagellum or to the background. A cell either divides or does not. Standard pipelines usually solve these discrete choices with local post-processing. Detection-based models such as YOLO^1^, Mask R-CNN ^2^, or StarDist ^3^ produce dense candidate outputs that are pruned by non-max suppression, while flow-field or semantic segmenters such as Cellpose ^4–7^, Omnipose ^8^, Mesmer ^9^, or nnU-Net ^10^ rely on watershed, connected components, or flow-based clustering to separate touching instances. Thresholds decide whether a candidate is accepted. For video analysis, linkers connect objects across frames to minimize distance-based objectives, e.g. linear assignment^11–16^, global network-flow or Viterbi assignment ^17–19^, or by training specialized learned trackers^20^. Division events are typically added by separate rules or by dedicated lineage-tracing methods^21–26^.

Doing segmentation and tracking in separate consecutive stages is practical, but each stage commits to a hard local decision before the next stage runs, so image evidence and temporal evidence are never weighed against each other in a single objective. A detection dropped at non-max suppression cannot be recovered by a temporally consistent lineage, and a link ruled out by the tracker cannot be reopened by a strong image cue two frames later. However, the optimal decision in one stage are intrinsically coupled to decisions in other stages. Such global discrete optimization problems are routinely solved by modern integer linear programming (ILP) solvers, which have become highly performant^27^. Here, we show that globally coupled video tracking problems in biology can be recast as integer programming problems, whose solutions can be efficiently found using modern ILP solvers. We present this as discrete inverse rendering: rather than forcing a detector to output a single answer per frame, we keep a large pool of candidate objects and let a solver select the subset that gives a globally consistent explanation of the video. The approach rests on two requirements shared by many biological tracking problems: the selected objects must reconstruct the image data (so the hypotheses are consistent with the pixels), and they must be temporally consistent (so the resulting tracks are biologically feasible). We showcase this in three distinct cases: suppression, path selection, and event-structured tracking.

## 2 Results

### 2.1 Framework

We consider an observed video *I*_1:*T*_ . Using any desired method, a set of candidate hypotheses is generated for each frame: splines, curve fragments or cell masks, with parameters *θ*_*i*_ and a renderer *R*(*θ*_*i*_) that predicts how each hypothesis appears in the image. We then introduce binary variables *x*_*i*_ ∈ {0, 1} that indicate hypothesis selection, with additional binary variables that encode e.g. temporal links (*y*_*ij*_), births (*b*_*i*_), deaths (*d*_*i*_), or division events (*s*_*i*_) when needed. We consider a general objective that balances an inverse rendering loss and a regularization loss:

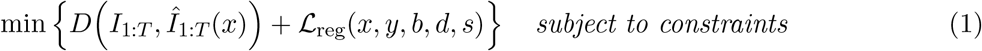

Here, *Î*_1:*T*_ (*x*) is the video reconstructed from the selected hypotheses, *D* is a discrepancy between the observed and reconstructed videos (image evidence), ℒ’_reg_ collects temporal-consistency and event penalties.

The objective above is general, but reduces to concrete integer programs whose forms depend on the problems. As a concrete example, consider tracking a fixed number of non-dividing cells: the reconstruction term is linear in the selection variables and no births, deaths, or divisions can occur, so the integer program reduces to

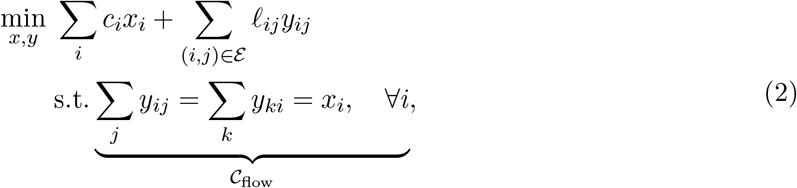

where *c*_*i*_ is candidate *i*’s reconstruction gain relative to the background and *ℓ*_*ij*_ is a temporal-consistency cost of linking candidates *i* and *j* across consecutive frames. The flow constraint *C*_flow_ requires every selected candidate to have exactly one predecessor and one successor (selected from possible links E), so segmentation (*x*_*i*_) and linking (*y*_*ij*_) are decided jointly by one solver against one objective: a candidate cannot survive without a temporally consistent partner, and a link cannot survive without visible support in the reconstruction. The problem can then be solved to certified optimality with CP-SAT^27^. The three applications below extend this template with problem-specific constraint families.

### 2.2 Suppression

As our first example application, we consider suppression in the context of predictions from deep neural models. Many image-analysis pipelines generate several overlapping hypotheses for the same object. Standard non-max suppression resolves this greedily by keeping high-scoring candidates and discarding lower-scoring overlapping ones. Such methods use no reconstruction check and no temporal information, and while the suppression rule itself can be simple, densely overlapping shapes typically require either substantial detector training or a dedicated learned suppression stage. Deeptangle^28^, for instance, learns a latent-space encoder whose clustering yields the final detections.

Here, we demonstrate that the suppression in dense *C. elegans* spline tracking can also be achieved with discrete integer programming and without the need for supervised suppression training. A deep-learning model^28^ produces many candidate worm splines per frame [Fig. 1A]. Instead of applying standard non-max suppression, we keep the large candidate pool and formulate selection as an integer program. Each spline is rendered onto the image by placing Gaussian blobs along the spline, and assigned a reconstruction cost. Temporal links keep identities consistent across frames, scored by a dynamic-time-warping distance between splines [Fig. 1B], and births (*b*_*i*_) and deaths (*d*_*i*_) absorb worms entering or leaving the field of view. Concretely, the integer program becomes

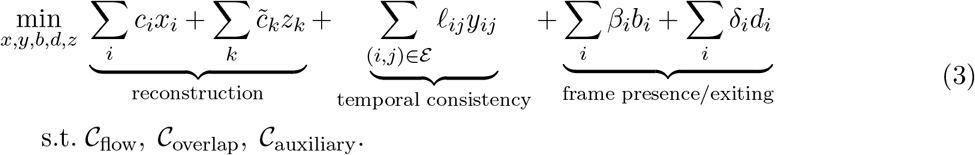

**Figure 1:**
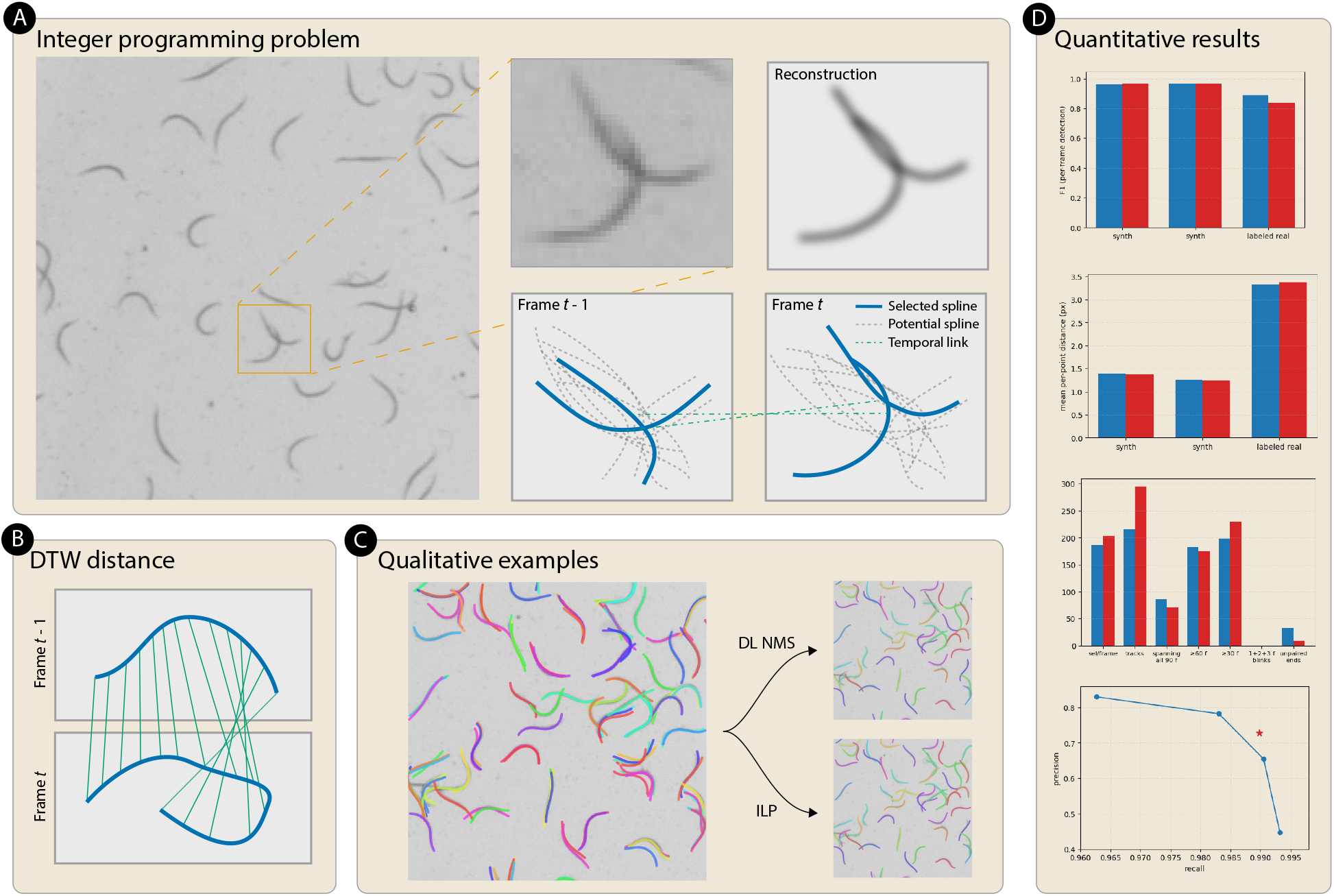
Suppression. **(A)** A deep network (deeptangle 28) proposes many overlapping worm splines per frame. Instead of non-max suppression we keep the whole pool and select a subset by integer programming: each spline is rendered as Gaussian blobs along its length, and the selected splines must reconstruct the image while temporal links keep identities consistent across frames. **(B)** Temporal consistency between consecutive frames is scored by a dynamic-time-warping distance between splines. **(C)** On dense C. elegans data, ILP selection and deep-learning non-max suppression yield visually comparable tracks. **(D)** Detection F1, mean per-point distance, track counts, and the precision–recall operating point on two synthetic densities (100 and 200 worms/frame) and hand-labelled real sequences: ILP selection from the raw candidate pool (red) matches deeptangle’s own trained suppression head (blue) across all metrics, reaching F1 = 0.89 and mean per-point distance 3.3 px on the real data. Precision-recall trade-off can be adjusted in deeptangle’s learned suppression algorithm, yet our default ILP algorithms improves on the Pareto front (bottom figure).

Here, *C*_flow_ represents the same constraints as in Eq. (2), but with the addition of the birth and death variables. For instance, the predecessor constraint becomes *b*_*i*_ + Σ_*j*_ *y*_*ji*_ = *x*_*i*_: a selected spline either appears through a birth (cost *β*_*i*_) or is linked to a predecessor in the previous frame, and analogously for the successor constraint with deaths (cost *δ*_*i*_). The overlap constraint *C*_overlap_ forbids pairs of near-identical splines (mask overlap ≥ 90%). Weaker overlaps that survive *C*_overlap_ make the reconstruction cost nonlinear in *x*, because pixels covered by more than one selected spline would otherwise be double-counted. Auxiliary variables *z*_*k*_, tied by *C*_auxiliary_ to the joint selection of an overlapping candidate set, carry correction terms 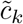 that cancel this double-counting.

In contrast to standard non-max suppression, which prunes greedily by score, this is an instance of discrete inverse rendering: choose the subset of renderings that best explains the image sequence. Because the criterion is image evidence rather than detector confidence, deeptangle’s learned suppression head is not required, and ILP tracks are visually comparable to deeptangle’s (Fig. 1C) and match its detection *F*_1_ across synthetic and real data (Fig. 1D).

### 2.3 Path selection

Our second motif is path selection. Some biological structures – flagella, axons, filaments, hyphae – are single connected curves with exactly two endpoints rather than sets of isolated objects. Local detectors (thresholding, skeletonisation, filament-response filters) readily produce candidate fragments [Fig. 2D], but many belong to background clutter rather than the target curve, and only the curve as a whole identifies which fragments are real. Integer programming provides an opportunity to give the optimization framework access to this key global information that is not available at the local scale: The selected path fragments must constitute a single connected path with exactly two endpoints and no branch points. We demonstrate this idea for path selection with flagellum tracking in a phase-contrast video of a swimming sperm cell, where moving microspheres produce local intensity profiles that mimic short flagellum segments. Frame by frame, a local detector cannot tell the two apart, and neural methods are typically needed to track such flagella^29,30^.

**Figure 2:**
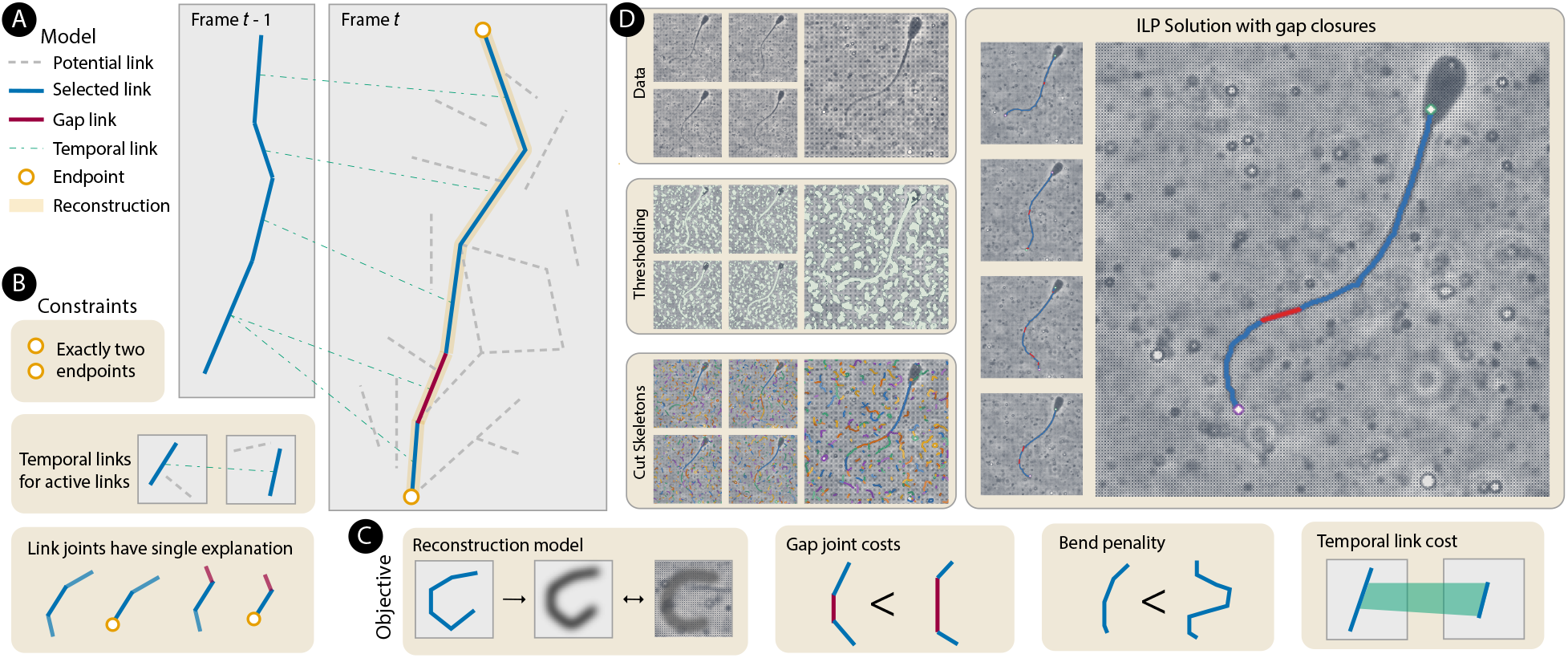
Path selection. **(A)** A flagellum is represented as a path built from short curve fragments. Candidate fragments and the gaps between them are rendered as Gaussian blobs, and the integer program selects one continuous path (blue), closing gaps where needed (red) and linking the path across frames (green). **(B)** Constraints enforce a valid path: exactly two endpoints, a single consistent explanation at each joint, and temporal links only between active fragments. **(C)** The objective combines a reconstruction term with penalties for gap joints, bending, and temporal change. **(D)** In phase-contrast video with distracting microspheres, thresholding and skeletonization yield many spurious fragments; the global program extracts the single flagellum with gap closures where frame-by-frame decisions fail. Video from29,30 .

The main hypothesis variables of the resulting integer program select fragments and, where local detection breaks, short closure segments, subject to a constraint that the selection forms one continuous path (Fig. 2A–B). Each selected fragment or closure contributes Gaussian blobs to the reconstructed image, and the objective balances reconstruction fidelity against penalties for gap closures, path curvature, and inter-frame change (Fig. 2C). The program takes the form

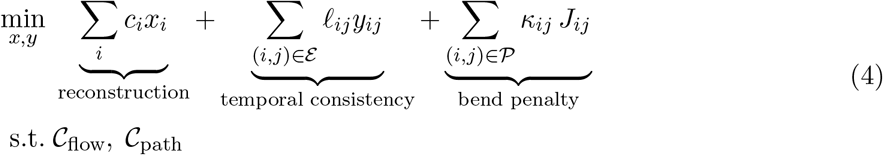

where *x*_*i*_ selects either a candidate fragment or a short gap-closure segment, *c*_*i*_ is its reconstruction cost (including a per-fragment activation cost for gap closures), *ℓ*_*ij*_ is the temporal cost linking selected segments across consecutive frames, and each *J*_*ij*_ is a binary variable representing a specific candidate join between two fragments *i* and *j* (enumerated at problem-build time, one per plausible pair), carrying its own precomputed bend penalty *κ*_*ij*_. *C*_flow_ enforces per-fragment temporal-link balance [Fig. 2A], while *C*_path_ enforces a single connected curve with exactly two endpoints [Fig. 2B]. In implementation, *C*_path_ combines per-fragment endpoint-flag variables with auxiliary topological-order variables that prevent cycles in the selected subgraph.

The resulting track [Fig. 2D] shows that the global constraints let a classical high-recall computer-vision pipeline track a flagellum in a cluttered environment. The hard decision – which fragments are flagellum and which are microsphere – is moved into the global objective, and the same construction would work with a DL detector front-end. This is a simple example, but the idea can readily be extended to more complex systems. Extension to multi-flagellar organisms (e.g. *Chlamydomonas*) requires relaxing the two-endpoint constraint, and extension to branched networks would require relaxing the no-branch condition.

### 2.4 Event-structured tracking

Our third example is event-structured tracking. In cell tracking^31–33^, the discrete decisions include object existence, temporal identity, birth, death, and division. A cell mask may be plausible in one frame but inconsistent over time. Conversely, a temporally plausible track may be unsupported by the image data.

We generate candidate cell masks by running Cellpose ^4^ on *K* image augmentations of each frame (identity plus perturbations in gamma, contrast, and brightness) and pooling all detected masks (Fig. 3A). Each detected mask is rendered as the raw pixels inside its footprint against a fixed background. The solver selects masks and lineage events jointly: reconstruction loss favours masks that explain the image, temporal consistency favors stable locations, shapes and areas, while division constraints enforce valid mother–daughter relationships (Fig. 3B). The full integer program is

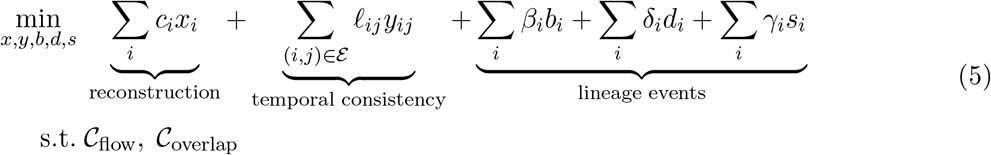

where *c*_*i*_ is the reconstruction cost of a candidate mask against the background, *ℓ*_*ij*_ is the temporal-consistency cost on linking cells *i* and *j* across consecutive frames, and *β*_*i*_, *δ*_*i*_, *γ*_*i*_ are the per-instance birth, death, and division penalties. The overlap constraint *C*_overlap_ is a hard exclusion here: no two selected masks may share a pixel, which makes the reconstruction cost stay linear in *x* and no auxiliary variables are needed. The flow constraint extends the simple form to a lineage graph: on entry, a selected cell *i* is either a fresh birth (*b*_*i*_ = 1) or has one predecessor via some *y*_*ji*_. On exit, it either dies (*d*_*i*_ = 1), continues via one successor *y*_*ij*_, or divides (*s*_*i*_ = 1, which increases the outgoing-link budget from one to two).

**Figure 3:**
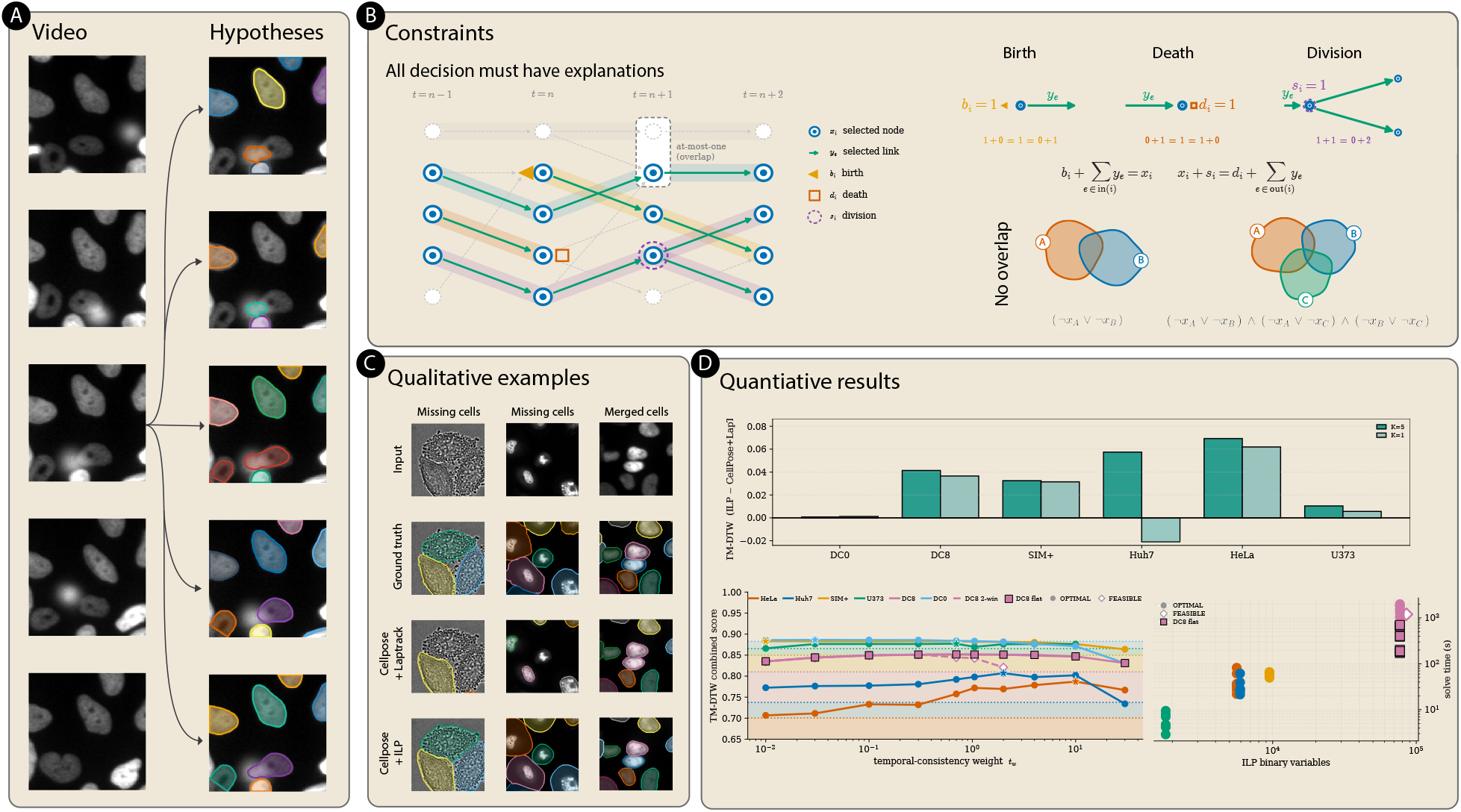
Event-structured tracking. **(A)** From each frame of a nuclei video we generate a pool of candidate cell masks. **(B)** Selection is posed as an integer program in which every selected mask, temporal link, birth, death, and division must be jointly explained: flow constraints tie each cell to a valid predecessor and successor, births and deaths absorb the ends of tracks, divisions enforce mother–daughter relationships, and overlapping masks are mutually exclusive. **(C)** Against a perframe segmentation linked by a standard tracker, joint selection recovers cells that are missed in single frames and separates masks that were wrongly merged. **(D)** Combined topology–morphology (TM-DTW) score across six Cell Tracking Challenge and DeepCell datasets, with solve time growing with the number of binary variables.

We evaluate the resulting tracks with a topology–morphology score (TM-DTW): the harmonic mean of a topology *F*_1_ over centroid-matched cells and a morphology term (a dynamic-time-warping distance between the matched boundary contours, mapped to [0, 1]). A method scores highly only when it recovers both the right cells and their shapes. TM-DTW complements the Cell Tracking Challenge’s TRA metric ^34^, which scores lineage topology alone, and its SEG metric, which scores per-mask IoU but not detection recall’ jointly scoring topology and morphology exposes shape errors that leave the tracking graph unchanged but degrade downstream biological measurements.

Across six benchmark videos (Table 1), joint selection improves TM-DTW over both Cellpose+LapTrack and Ultrack on five of six sequences, with the largest gains where the per-frame segmentation is least reliable (Fig. 3C). On DIC-C2DH-HeLa, TM-DTW rises from 0.70 (Cellpose+LapTrack) and 0.71 (Ultrack) to 0.77’ on Fluo-C2DL-Huh7, detection *F*_1_ roughly doubles from 0.31 to 0.58 (both baselines) while TM-DTW rises from 0.74 to 0.80’ on the dense seq 08, detection *F*_1_ climbs from 0.79 to 0.92 and TM-DTW from 0.81 to 0.85. On U373 and seq 00 the three methods differ by less than 0.02 TM-DTW, as expected when per-frame segmentation contains little for the tracker to correct. We further perform a temporal-weight sweep by multiplying the temporal consistency and lineage events terms by a weight *t*_*w*_. Fig 3D shows a broad optimum near *t*_*w*_ ≈ 1–10. For instance, topology *F*_1_ on DIC-HeLa rises from 0.74 at *t*_*w*_ = 0.01 to 0.95 at *t*_*w*_ = 10, then collapses at *t*_*w*_ = 100 when lineage feasibility overrides the image (Fig. 3D). Solve times ranged from 7 s (U373) to 62 s (SIM+) at *t*_*w*_ = 1 on the four smaller sequences we timed, while the dense seq 08 was solved in ∼650 s.

**Table 1:** Segmentation-and-tracking performance across six Cell Tracking Challenge and DeepCell videos. Higher is better for both metrics. Best per row in **bold** (within 0.005 both marked).

| Dataset | Cellpose+LapTrack |  | Ultrack |  | Ours |  |
| --- | --- | --- | --- | --- | --- | --- |
| | TM-DTW | $F_1$ | TM-DTW | $F_1$ | TM-DTW | $F_1$ |
| DIC-C2DH-HeLa | 0.700 | 0.809 | 0.711 | 0.843 | **0.770** | **0.880** |
| Fluo-C2DL-Huh7 | 0.737 | 0.308 | 0.735 | 0.309 | **0.795** | **0.584** |
| Fluo-N2DH-SIM+ | 0.850 | 0.971 | 0.851 | 0.974 | **0.882** | **0.977** |
| PhC-C2DH-U373 | 0.866 | 0.925 | **0.883** | **0.949** | 0.876 | 0.937 |
| DeepCell seq_00 | **0.883** | 0.927 | **0.885** | **0.934** | **0.883** | 0.930 |
| DeepCell seq_08 | 0.810 | 0.794 | 0.810 | 0.795 | **0.851** | **0.923** |

This construction is closely related to Ultrack ^25,26^, which likewise selects cell hypotheses and lineage events with integer programming. The difference is what scores a hypothesis: rather than relying on tracking confidence alone, we score masks by how much their presence explains the observed image relative to the background estimate, so that image evidence and lineage feasibility are weighed against each other in a single objective. Reconstruction then acts as a data-driven regulariser: dropping masks that are temporally convenient but visually unimportant, and rescuing cells that a per-frame segmenter missed but that image evidence and a valid lineage together justify.

## 3 Discussion

We have presented discrete inverse rendering as a general framework for resolving hard combinatorial decisions that arise in biological video analysis. Instead of committing to one detection, segmentation, or local structure before temporal and structural evidence is considered, the method retains a pool of candidate renderings and selects a globally consistent explanation of the image sequence. Across dense worm detections, fragmented sperm flagella, and dividing-cell lineages, this approach matched specialised pipelines.

Each of the three example applications specializes the framework defined by Eq. (1). In each case, candidate hypotheses are rendered, their agreement with the image data is converted into reconstruction costs, and binary variables select a feasible subset under global consistency constraints. The instantiated variables, regularisation terms, and feasible sets depend on the biological problem: spline suppression requires auxiliary variables that correct the reconstruction cost in overlapping regions, path selection requires connectivity, endpoint, and bending terms, and cell tracking requires lineage-event variables (births, deaths, divisions).

The framework separates candidate generation from global interpretation. The candidate pool may be produced by a neural detector, a classical image-processing pipeline, perturbations of a segmenter, or another high-recall proposal mechanism. Domain knowledge then enters through explicit constraints and penalties, while the raw image provides a common evidence term through reconstruction. This makes object existence, temporal identity, and structural validity mutually informative rather than sequential decisions. Integer-programming approaches to lineage reconstruction, including Ultrack ^25,26^, provide an important precedent for this strategy in cell tracking. The present results broaden that perspective and show that reconstruction-based scoring is particularly useful when confidence from a per-frame predictor is insufficient.

Solving discrete optimization problems is harder than smooth, continuous optimization, and the present approach is only feasible due to modern advances in integer programming ^27^. Runtime remains a concern, and it does not scale with a single quantity: it depends on the number of candidate hypotheses, the density of overlap and temporal edges, the number of event and auxiliary variables, and the ambiguity of the resulting optimisation problem. The reported problems reached certified optimality in seconds to minutes, with the densest benchmark (DeepCell seq 08, ∼200 cells per frame across 71 frames) taking ∼650 s on commodity hardware. For longer recordings, overlapping temporal windows provide a practical decomposition: part of each overlap can be fixed to preserve identities across a seam, while the remaining frames are re-optimised using newly available evidence. With a fixed window size, this makes computational cost grow approximately linearly with video length, but it forfeits a certificate of global optimality for the complete video and may introduce seam-dependent decisions. The method is also bounded by the candidate pool: a missing hypothesis cannot be recovered, and the present linearised objective selects among fixed geometries rather than refining them continuously.

A major benefit of inverse rendering formulations is *auditability*. The reconstructed video and the contribution of each selected candidate can be regenerated from the input, candidate pool, and recorded solution, while the objective terms and active constraints expose how image evidence, temporal consistency, and lineage events were weighed. Acceptance or rejection is nevertheless a global property of the solution: a hypothesis may be excluded because it conflicts with a better combination of alternatives, not simply because one of its individual costs is poor. Certified optimality establishes that no feasible solution has a lower objective cost.

Differentiable inverse rendering^35^ addresses the complementary continuous problem: refining shape, pose, or appearance parameters when the scene structure is fixed. It does not directly provide certified hard combinatorial selection, whereas the integer formulation used here makes those choices but keeps the geometry of each candidate fixed. The two can be combined naturally. For instance, differentiable inverse rendering can be used for sub-pixel spline refinement of the discrete output of nematode tracking ^36^.

## Supporting information

Ablations, confidence intervals and additional metrics

## Acknowledgements

This work was supported by the Novo Nordisk Foundation under grants NNF20OC0062047 and NNF25OC0106165.

## Author contributions

J.B.K. conceived the framework and its application to each of the three motifs. F.Z. implemented the integer-programming models, benchmarking pipeline, and evaluation. J.B.K. supervised the project. Both authors wrote the manuscript.

## STAR Methods

### Key resources table

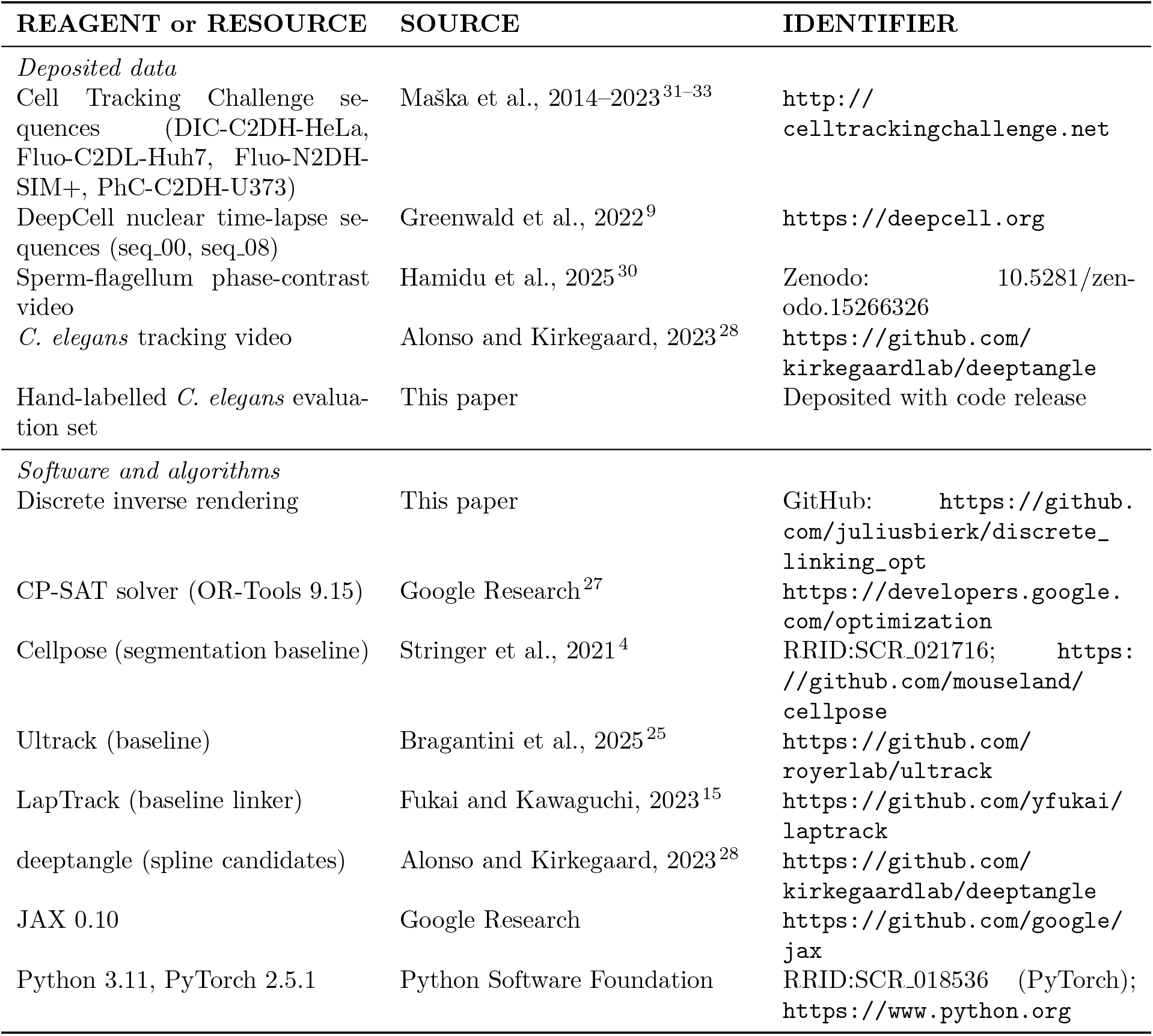

## Resource availability

### Lead contact

Requests for further information should be directed to Julius B. Kirkegaard.

### Materials availability

This study did not generate new physical materials.

### Data and code availability

- All benchmark datasets used in this study are publicly available through the sources listed in the Key resources table.
- All original code has been deposited at https://github.com/juliusbierk/discrete_linking_opt and is publicly available.

## Method details

### Constraints

Overlap: for any pair (*i, j*) whose masks overlap beyond a modality-specific threshold (90% area for splines, any pixel for cells), *x*_*i*_ + *x*_*j*_ ≤ 1. Flow: for every selected candidate *x*_*i*_ = 1 in frame *t < T*, exactly one of {*y*_*ij*_ : *j* ∈ ℰ_*i*_} ∪ {*d*_*i*_} activates, and symmetrically for predecessors. Path (flagella): exactly two vertices have degree 1 and all others degree 2 in the selected subgraph. Division: a split indicator *s*_*i*_ ≤ *x*_*i*_ modifies the flow balance at node *i* to *d*_*i*_ + ∑ _*e*∈out(*i*)_ *y*_*e*_ = *x*_*i*_ +*s*_*i*_, so *s*_*i*_ = 1 forces *i* to have two outgoing links (its daughters), which are then determined by which two link variables are jointly selected’ *s*_*i*_ is fixed to zero when *i* has fewer than two candidate successors. Auxiliary: for each auxiliary variable *z*_*k*_ representing the joint selection of candidates (*i*_1_, …, *i*_*m*_), the linearisation 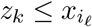 for each *ℓ* and 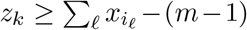 enforces 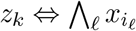 so the correction 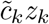 activates exactly when the corresponding candidates are jointly selected.

### Candidate generation

Splines come from a trained deeptangle network ^28^ at its lowest score threshold. Flagellum fragments come from local thresholding and morphological skeletonisation with gap-closure candidates enumerated between endpoints. Cell masks come from Cellpose ^4–7^ run on *K* image augmentations of each frame (identity plus gamma 0.5, gamma 2.0, contrast 1.4, and brightness −0.10) and pooled; increasing *K* enlarges the pool without changing the objective (K-ablation, Fig. S1 in the Supplementary Information).

### Solver settings

OR-Tools CP-SAT 9.15 with default parameters. The DeepCell seq 08 *t*_*w*_-sweep (Fig. 3D) is solved via the windowed decomposition below and returns near-optimal FEASIBLE per window at the 600 s cap. The sweep covers *t*_*w*_ ∈ {0.01, 0.03, 0.1, 0.3, 0.7, 1, 2, 4, 10, 30, 100}.

### Windowed decomposition

For videos too long to solve monolithically, the driver splits the video into overlapping windows of *W* frames, adjacent windows sharing *H* frames of overlap. The first *F* ≤ *H* frames of each overlap are hard-frozen from the previous window’s solution, so cell identities agree across the seam; the remaining *H* −*F* frames give the next window room to revise its committed decisions in light of new evidence. Each window’s ILP is built and solved independently, and per-window tracks are stitched into one global lineage by matching selected renderings across the overlap. For the DeepCell seq 08 *t*_*w*_-sweep we used *W* = 40, *H* = 8, *F* = 4 (two windows covering the 71-frame sequence) with a 600 s cap per window.

### Quantification and statistical analysis

#### Detection

*F*_1_. For each frame, predicted cell centroids are matched to ground-truth centroids by Hungarian assignment on Euclidean distance, capped at a per-dataset radius (80 px for DIC-C2DH-HeLa and PhC-C2DH-U373, 50 px for Fluo-C2DL-Huh7, 30 px for Fluo-N2DH-SIM+, and 15 px for both DeepCell sequences, chosen on the order of a typical cell radius). The Hungarian match yields true positives (matched pairs), false negatives (unmatched ground-truth cells), and false positives (unmatched predictions); the topology score is their *F*_1_:

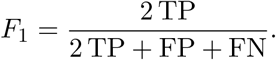

#### TM-DTW score

Using the same centroid matches as the detection *F*_1_ above, each matched pair contributes a morphology term 1 − DTW(*c*_pred_, *c*_gt_)*/L*_max_, where *c*_pred_ and *c*_gt_ are boundary contours sampled at fixed arc-length spacing, DTW is the dynamic-time-warping distance, and *L*_max_ normalises to [0, 1]. The per-sequence morphology score *M* is the mean over matched pairs. TM-DTW is the harmonic mean 2*F*_1_*M/*(*F*_1_ + *M*) of the detection *F*_1_ and *M*, so both topology and shape must be recovered to score highly.

#### Confidence intervals

Per-frame TM-DTW values are bootstrapped (5000 resamples over frames) to give 95% confidence intervals; the full CI table is in Table S1 of the Supplementary Information. For the sequences in Table 1, 95% CI half-widths range from ±0.001 for DeepCell seq 08 (71 frames) to ±0.04 for DIC-C2DH-HeLa (7 frames), so ranking differences at the 0.05 level are not sensitive to individual frames even on the shortest sequences.

#### Baselines

Cellpose+LapTrack uses Cellpose 3^6^ with default cyto3 weights per frame, then links masks across frames with LapTrack ^15^, a linear-assignment particle tracker. Search radius scales with the median cell diameter. On the dense DeepCell seq 08 (which has many divisions) LapTrack’s splitting mode is enabled with a 120-pixel cutoff, while the other five sequences use LapTrack without splitting. Ultrack uses each dataset’s default configuration file from the Ultrack repository. Hypotheses are its own Cellpose ensembles. For DIC-C2DH-HeLa and Fluo-N2DH-SIM+, TRA and DET scores from the Cell Tracking Challenge evaluator ^34^ are reported in Table S2 of the Supplementary Information.

