## Supplementary material for "Discrete Inverse Rendering: Biological Image Analysis with Integer Programming": Ablations, confidence intervals and additional metrics

### Supplementary Information for "Discrete Inverse Rendering: Biological Image Analysis with Integer Programming"

#### S1. Candidate-pool ablation

The reconstruction-scored objective is unchanged by the size of the candidate pool: increasing the number of Cellpose image augmentations $K$ increases the number of binary variables in the integer program but does not degrade tracking quality. Figure S1 shows TM-DTW and integer-variable count for $K \in \{1, 2, 3, 5\}$ on two Cell Tracking Challenge sequences. TM-DTW rises modestly and plateaus by $K = 3$ on both sequences, while the number of binary variables grows roughly linearly with $K$. All runs reached CP-SAT OPTIMAL.

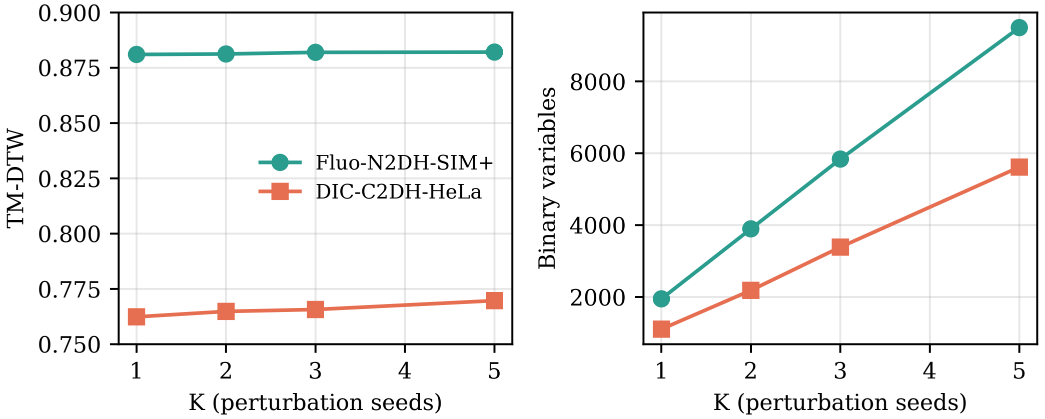


Figure S1: **Candidate-pool ablation ($K$-sweep).** Effect of increasing the number of Cellpose image augmentations $K$ on tracking quality (left, TM-DTW) and problem size (right, number of binary variables). TM-DTW improves modestly with $K$ and plateaus by $K = 3$; the number of binary variables grows roughly linearly with $K$.

#### S2. Bootstrapped confidence intervals for TM-DTW

Per-frame TM-DTW values were bootstrapped with 5,000 resamples over frames to give 95% percentile confidence intervals for the point estimates reported in Table 1 of the main text. Table S1 summarises. Half-widths range from ±0.001 for DeepCell seq_08 (71 frames) to ±0.04 for DIC-C2DH-HeLa (7 frames), so the ranking differences reported in the main text are not sensitive to individual frames.

Table S1: Bootstrapped 95% confidence intervals for TM-DTW (5,000 resamples over frames).

| Dataset | Ours: TM-DTW [95% CI] | Cellpose+LapTrack: TM-DTW [95% CI] |
|---|---|---|
| DIC-C2DH-HeLa | 0.770 [0.740, 0.800] | 0.700 [0.664, 0.730] |
| Fluo-C2DL-Huh7 | 0.795 [0.778, 0.812] | 0.737 [0.723, 0.752] |
| Fluo-N2DH-SIM+ | 0.882 [0.878, 0.886] | 0.850 [0.844, 0.855] |
| PhC-C2DH-U373 | 0.876 [0.855, 0.891] | 0.866 [0.838, 0.892] |
| DeepCell seq_00 | 0.883 [0.880, 0.886] | 0.883 [0.879, 0.886] |
| DeepCell seq_08 | 0.851 [0.850, 0.853] | 0.810 [0.807, 0.813] |

#### S3. Cell Tracking Challenge evaluator scores (TRA, DET)

For the two Cell Tracking Challenge sequences on which the official evaluator was run, Table S2 reports TRA (tracking accuracy) and DET (detection accuracy) alongside those of the Cellpose+LapTrack baseline. TRA and DET are the graph-based metrics defined in the Cell Tracking Challenge [31 in main]; higher is better on both. On DIC-C2DH-HeLa the reconstruction-scored variant improves TRA by $\sim$0.02 over the baseline; on Fluo-N2DH-SIM+ both approaches are within 0.005 TRA of each other, reflecting the easier per-frame segmentation on this sequence.

Table S2: Cell Tracking Challenge evaluator scores (TRA, DET).

| | Ours | | Cellpose+LapTrack | |
|---|---|---|---|---|
| Dataset | TRA | DET | TRA | DET |
| DIC-C2DH-HeLa | 0.979 | 0.977 | 0.958 | 0.960 |
| Fluo-N2DH-SIM+ | 0.979 | 0.982 | 0.975 | 0.978 |
